# CLEAN AND FOLDED: OPTIMIZED PRODUCTION OF HIGH QUALITY RECOMBINANT HUMAN INTERFERON-Λ1

**DOI:** 10.1101/2020.12.09.417527

**Authors:** Aram Shaldzhyan, Nikita Yolshin, Yana Zabrodskaya, Tatiana Kudling, Alexey Lozhkov, Marina Plotnikova, Edward Ramsay, Aleksandr Taraskin, Peter Nekrasov, Mikhail Grudinin, Andrey Vasin

## Abstract

Type III interferons exhibit antiviral activity against influenza viruses, coronaviruses, rotaviruses, and others. In addition, this type of interferon theoretically has therapeutic advantages, in comparison with type I interferons, due to its ability to activate a narrower group of genes in a relatively small group of target cells. Hence, it can elicit more targeted antiviral or immunomodulatory responses. Obtaining biologically-active interferon lambda (hIFN-λ_1_) is fraught with difficulties at the stage of expression in soluble form or, in the case of expression in the form of inclusion bodies, at the stage of refolding. In this work, hIFN-λ_1_ was expressed in the form of inclusion bodies, and a simple, effective refolding method was developed. Efficient and scalable methods for chromatographic purification of recombinant hIFN-λ_1_ were also developed. High-yield, high-purity product was obtained through optimization of several processes including: recombinant protein expression; metal affinity chromatography; cation exchange chromatography; and an intermediate protein refolding stage. The obtained protein was shown to have expected, specific biological activity in line with published effects: induction of MxA gene expression in A549 cells.

## 1. INTRODUCTION

Interferons play a critical role in the immune response by suppressing viral spread in the early stages of infection and form the first line of defense against viral infection in mammals [1,2]. According to amino acid sequence and type of receptor through which signal transmission is mediated, interferons are divided into three groups: types I, II, and III [1,3]. Type III interferons (lambda interferons, IFN-λ) are related to type I interferons (interferon-α and interferon-β) and demonstrate a similar antiviral effect [4,5]. Type III interferons are promising therapeutic agents that can induce a more targeted antiviral or immunomodulatory response than other interferon types [6,7]. Such interferons primarily affect viruses that target cells of the respiratory tract, gastrointestinal tract, or liver [8]. Thus, type III interferons exhibit antiviral activity against a number of respiratory and gastrointestinal viruses: influenza virus; metapneumovirus; respiratory syncytial virus; coronavirus; and rotavirus [6]. Availability of a simple, effective method for producing type III interferons is a necessary prerequisite for further research of its potential as an antiviral or immunomodulatory agent. In previously published work, methods for production of recombinant hIFN-λ_1_ in soluble form with the proteolytically cleaved S-tag [9], and in the form of inclusion bodies with 6His-tag [5,10], have been proposed. In this work, we cloned and expressed recombinant human interferon-λ_1_ (hIFN-λ_1_). Along with the 6His-tag, inclusion body approach, we propose: a simple and effective refolding method; and an efficient, potentially-scalable chromatographic purification method. The hIFN-λ_1_ produced features specific activity suitable for research and, potentially, future diagnostic or therapeutic application.

## 2. MATERIAL AND METHODS

### 2.1. Materials

Reagents obtained from Sigma included: tris base; Na_2_HPO_4_; NaCl; EDTA; PMSF; Tween 20; glycerol; acetonitrile; TFA; trehalose; L-arginine; sodium acetate; acetic acid; HCl; and urea. HisTrap FF Crude and Source 15S 4.6/100 columns were obtained from GE Healthcare; BioScale Mini P6 cartridges and TGX anyKD precast SDS-PAGE gels were obtained from Bio-Rad. For endotoxin evaluation, the LAL Chromogenic Endpoint Assay (#HIT302, Hycult Biotech) was used. For enzymatic hydrolyses, trypsin and chymotrypsin (Promega) were used. Matrices for MALDI-TOF mass spectrometry (HCCA) were manufactured by Bruker.

Lambda interferon levels were evaluated by human IL-29/IL-28B (IFN-lambda 1/3) DuoSet ELISA (DY1598B, R&D Systems, USA). Biological activity experiments were performed using: the A549 cell line (ATCC, CCL-185); F12K media, fetal bovine serum, and DPBS manufactured by Gibco (USA); and 12-well plates from Thermo Scientific Nunc (USA). Total RNA isolation and realtime PCR were performed using: TRIZol reagent (Invitrogen, USA); M-MLV reverse transcriptase, MMLV Reaction Buffer, and ultrapure water (Promega, USA); BioMaster HS-qPCR (BioLab-Mix, Russia); and oligo(dT)16 primers (DNA-Synthesis, Russia).

### 2.2. Plasmid construction

*In silico* optimized sequence (https://www.genscript.com/tools/gensmart-co-don-optimization), encoding hIFN-λ_1_ #Q8IU54 [11], was synthesized by Evrogen and cloned into the pet302-NT-His plasmid. That vector contains a sequence encoding a polyhistidine tag at the Nterminus of synthesized protein. The resultant construct (pet302-NT-His-hIFN-λ_1_) was checked for errors by sequencing.

### 2.3. Protein expression

*Escherichia coli* cultures (BL-21 DE3) were transformed with pet302-NT-His-hIFN-λ_1_ and seeded on Luria Bertani (LB) agar plates containing ampicillin (100 μg/ml). Individual colonies were picked and grown overnight in LB media (0.5% yeast extract, 1% Bacto tryptone, 1% NaCl) containing ampicillin. The overnight cultures were then used to inoculate fresh LB medium containing ampicillin (OD_600_ = 0.1 at start of new incubation). Bacterial cultures were cultivated with agitation (orbital shaker) to an OD_600_ value of 0.7. To optimize cultivation conditions, protein expression was induced by various IPTG concentrations (0.01, 0.05, 0.1 mM), and cultures were incubated further (orbital shaker, 32°C or 37°C, 3 hours). Cultures were sonicated (MSE ultrasonic disintegrator) on ice (10 pulses, of 10 seconds each, in 30-second intervals), followed by centrifugation for 1 hour (20,000 g at ^+^4°C). Supernatants and pellets were collected separately. Levels of hIFNλ_1_ expression, and its presence in supernatants or pellets (inclusion bodies), were determined by polyacrylamide gel electrophoresis (SDS-PAGE).

### 2.4. Protein purification and refolding

Chromatographic purification was performed on a GE Healthcare AKTA pure 25M system. Primary purification of hIFN-λ_1_ was performed by immobilized metal affinity chromatography (IMAC) under denaturing conditions. Cells were precipitated by centrifugation for 10 min at 4,500 g (^+^4°C). Next, cell pellets (1 g wet weight) were resuspended in 30 ml of denaturing buffer (20 mM sodium phosphate, 300 mM sodium chloride, 8 M urea, 20 mM imidazole, pH 7.8, 1 mM PMSF); suspensions were sonicated (MSE ultrasonic disintegrator) using 10 pulses (30 seconds each, at 2 minute intervals) on ice. Lysate was clarified by centrifugation for 20 minutes at 13,000 g (^+^10°C).

A 1 mL HisTrap FF Crude column was equilibrated with 5 ml of binding buffer (30 mM sodium phosphate, 300 mM sodium chloride, 8 M urea, 20 mM imidazole, pH 7.8) at a flow rate 0.5 ml/min. Clarified lysate was loaded onto the column (at a flow rate of 0.2 ml/min), followed by washing with 20 ml of binding buffer. The target protein was eluted with 5 ml of elution buffer (30 mM sodium phosphate, 300 mM sodium chloride, 8 M urea, 500 mM imidazole, pH 7.8) at a flow rate of 0.2 ml/min. Eluate was monitored by optical density (280 nm wavelength), and fractions of the target protein with an OD_280_ greater than 300 mAU were collected. EDTA (0.5 M) was added to target protein fractions to a final concentration of 5 mM. Protein concentration was determined by chromatogram integration using Unicorn 6 software, assuming: the absorbance of a 0.1% hIFN-λ_1_ solution at 280 nm to be 0.85 per cm of optical path (calculated based on primary protein sequence using the ProtParam service).

Protein refolding was performed by the dilution method. Chilled refolding solution was placed on a magnetic stirrer, and a solution of denatured protein (2 mg/ml) was added dropwise to a final concentration of 0.1 mg/ml. The resulting solution was stirred for 10 minutes and then incubated for 24 hours at ^+^4°C. The resulting refolded protein solution was filtered through a PES syringe filter (0.45 μm pore size).

Final purification was performed by cation exchange chromatography. A SOURCE™ 15S 4.6/100 column was equilibrated with 8 ml of start buffer (20 mM Tris hydrochloride, 2 mM EDTA, pH 7.5) at a flow rate of 2 ml/min. Next, refolded and filtered protein solution was loaded onto the column, followed by column washing with 10 ml of start buffer (2 ml/min). Target protein was eluted with a linear salt gradient (0 to 1 M NaCl) in start buffer at a flow rate of 2 ml/min for 15 minutes. Eluate was monitored by optical density at 280 nm; fractions (0.5 ml) featuring OD_280_ values above 25 mAU were collected. Fractions were analyzed by polyacrylamide gel electrophoresis. Fractions containing target protein with minimal impurities were combined. The Bio-Scale Mini P6 Desalting Cartridge (Bio-Rad) was used to transfer into phosphate buffered saline (pH 7.2). Protein concentration was determined by the Lowry method.

### 2.5. Product analysis

#### 2.5.1 SDS-PAGE

Purified protein was analyzed by SDS-PAGE using the Laemmli method under reducing conditions [12]. Electrophoresis materials used (Bio-Rad) included: ‘Any kD TGX’ precast gels; Kaleidoscope Plus Protein Ladder; PowerPac Basic DC power source; and Mini Protean Tetra cell. Gels were stained with a Coomassie solution as described [13]. Gels were visualized using the Chem-iDoc MP gel documentation system and analyzed using Bio-Rad Image Lab Software.

#### 2.5.2 Enzyme-linked immunosorbent assay

ELISA was carried out using commercial reagents. Levels of hIFN-λ_1_ were evaluated by human IL-29/IL-28B (IFN-lambda 1/3) DuoSet ELISA (DY1598B, R&D Systems, USA), according to manufacturer’s instructions. A different commercial kit, Mouse IL-28 A/B (IFN-lambda 2/3) DuoSet ELISA (DY1789B, R&D Systems), was used to assess cross-reactivity.

#### 2.5.3 MALDI-TOF

The amino acid sequence of recombinant 6xHis-hINF-λ_1_ protein was confirmed using MALDI-TOF mass spectrometry. Following PAGE, fragments were excised from gel and washed twice for dye removal (30 mM NH_4_HCO_3_ in 40% acetonitrile). Gel fragments were next dehydrated with 100% acetonitrile, dried in air, and subjected to enzymatic digestion by trypsin or chymotrypsin. Trypsin (20 ug/ml in 50 mM NH_4_HCO_3_) was added to gel, and incubation was performed for 2 hours at 60°C. Incubations with chymotrypsin (20 ug/ml in 100 mM Tris-HCl, 1 mM CaCl_2_, pH 8.0) were performed for 18 hours at 25°C. In both cases, reactions were stopped with 1% TFA in 10% acetonitrile. Peptides after enzymatic digestion were mixed with HCCA matrix, put onto the metal target, and their mass spectra recorded in positive ion registration reflective mode using the UltrafleXtreme MALDI-TOF mass spectrometer (Bruker). Spectra were processed in flexAnalysis software (Bruker). Protein identification was performed in BioTools (Bruker) using MASCOT (http://www.matrixscience.com/). It should be noted that: recombinant protein amino acid sequence (Fig. 1b) was added to the local database; during the identification process, the two databases (local and SwissProt, https://www.uniprot.org/) were used simultaneously. Mass tolerance was established as 20 ppm; oxidation of methionine was indicated as a variable modification. Identification was considered reliable based on spectrum score and threshold (p < 0.05).

**Figure 1.**
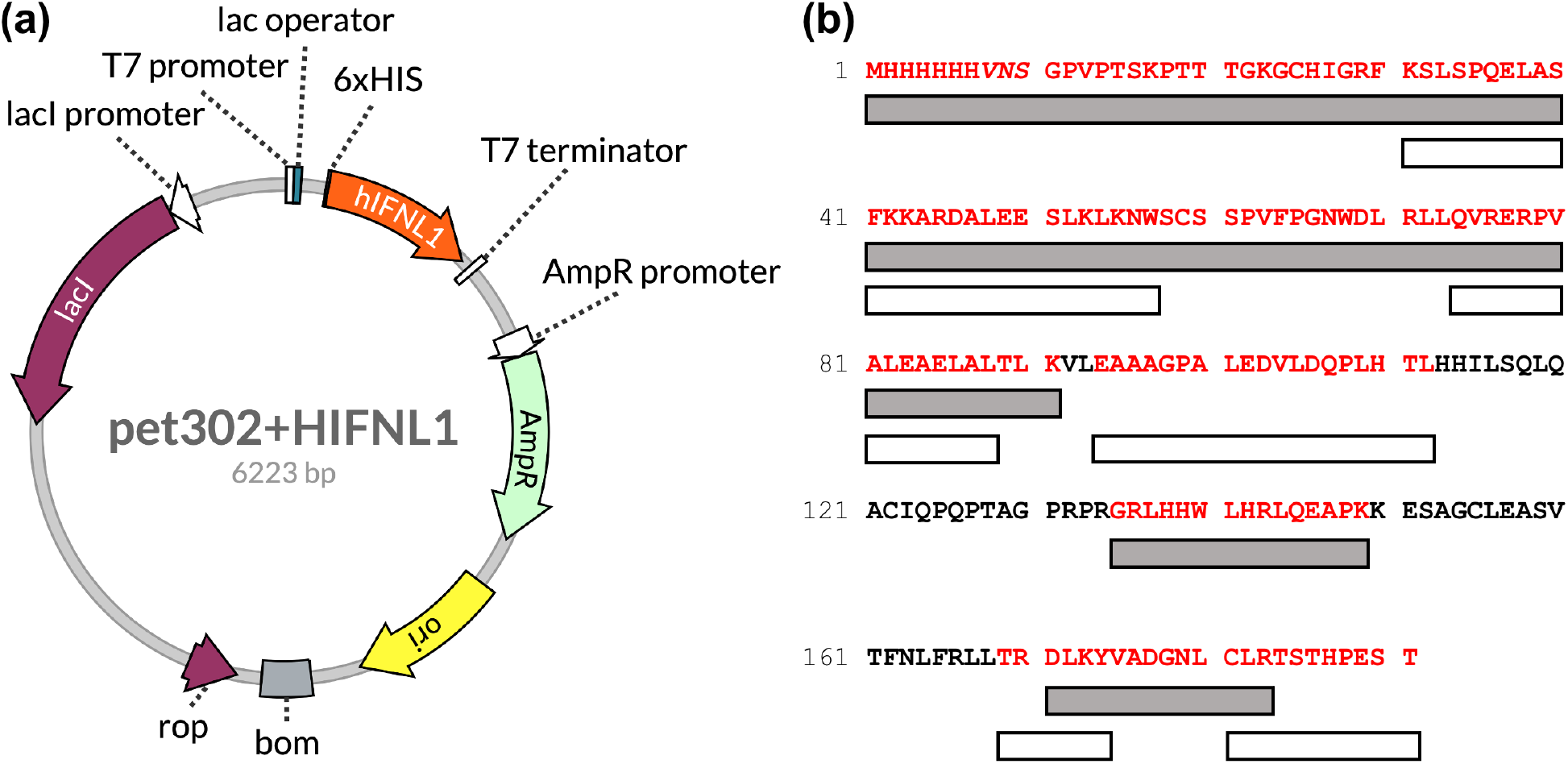
**(a)** Specifics of the pet302+ hIFN-λ_1_ expression plasmid. **(b)** Amino acid sequence of the 6xHis-hIFN-λ_1_ recombinant protein. Additional aa residues between the polystydine tag and hIFN-λ_1_ are indicated in italics. Sequence regions identified using mass spectrometry (section 3.4.1) are highlighted: black bars and white bars indicate enzymatic digestion with trypsin or chymotrypsin, respectively.

### 2.6. Biological activity

Type I and type III interferons induce the expression of ISGs (interferon stimulated genes) through their heterodimeric receptors. One of the key antiviral ISGs is MxA [3]. Unlike most ISGs, the Mx genes are not characterized by constitutive expression; their expression is specifically caused by the action of type I and III IFNs. This makes MxA an excellent marker of activation of an IFN-dependent response [14]. Thus, biologically active hIFN-λ_1_ should induce MxA expression. To determine changes in MxA mRNA level, real-time qPCR analysis was performed following sample preparation (RNA isolation, reverse transcription)

#### 2.6.1 Cell treatment

A549 (carcinomatous alveolar basal epithelial cell line) cells were used. Cells were cultured using: F12K media with 1% GlutaMax and 10% fetal bovine serum; and 12-well plates (5×10^5^ cells/well). Monolayers were washed with DPBS and treated with 10 ng/ml hIFN-λ_1_. After 10 h or 24 h of incubation (37°C, 5% CO_2_, humidification), hIFN-λ_1_ was removed. Cell monolayers were washed again with DPBS. The treated cell cultures and non-treated controls were harvested.

#### 2.6.2 RNA isolation

Total RNA was isolated from A549 cells using TRIZol reagent according to manufacturer’s instructions. RNA concentration and general integrity were measured using a NanoDrop ND-1000 spectrophotometer (NanoDrop Technologies, USA).

#### 2.6.3. Reverse transcription reaction

Following RNA extraction, samples were reverse transcribed using M-MLV reverse transcriptase (M-MLV RT) as described [15]. A mixture of 2 μg total RNA and 1 μg oligo(dT)_16_ primers, adjusted with ultrapure water to a final volume of 12 μl, was incubated at 70°C for 5 min. Tubes were immediately cooled on ice, and a mix was then added containing: 5 μl 5x MMLV Reaction Buffer; 1.5 μl 5 mM dNTPs; 1 μl M-MLV RT; and 5.5 μl ultrapure water. Complementary DNA synthesis was carried out at 42°C for 60 min. Enzyme (M-MLV RT) was inactivated by heating (65°C, 5 min). Products were stored at ^−^20°C.

#### 2.6.4. Real-time qPCR

Real-time qPCR assays were performed using the CFX96 Real-Time PCR System (Bio-Rad). Reaction mixtures contained: 12.5 μl of 2x BioMaster HS-qPCR; 0.5 μl (0.6 μM) of each primer and probe; 2 μl of first-strand cDNA; and 9 μl of ultrapure water. Amplification was performed using the following conditions: an activation step (95°C, 5 min); and 40 cycles of amplification (95°C for 15 sec; 60°C for 60 sec). Fluorescence detection was performed in each cycle at the 60°C step. The comparative cycle threshold method (ΔΔCT) was used to quantitate MxA expression. Results were normalized to GAPDH endogenous reference expression levels. At least three biological replicates were used for each data point. Statistical analyses were performed using GraphPad Prism 6.0.

## 3. RESULTS AND DISCUSSION

### 3.1. Plasmid construction

An *in silico* optimized (http://genomes.urv.es/OPTIMIZER/obtimized.php) sequence encoding hIFN-λ_1_ was synthesized and cloned into pet302 NT-His plasmid by Evrogen. The resultant plasmid was verified by sequencing and found to contain no errors. The pet302 NT-His vector contains sequence encoding a 6 histidine tag at the N-terminus of recombinant protein. The resultant recombinant protein amino acid sequence, encoded by the construct (plasmid pet302 + hIFN-λ_1_), is shown in Figure 1b.

### 3.2 Protein expression

Upon IPTG induction, a BL-21 DE3 cell culture transformed with the pet302-NT-His-hIFN-λ_1_ plasmid showed expression of a protein with a molecular weight of about 21 kDa. The theoretical molecular weight of the protein was 21.3 kDa. In the experiment, the concentration of inducer and the incubation temperature were varied. The incubation time with the inducer was 3 hours. Each sample was fractionated (Figure 2a).

**Figure 2.**
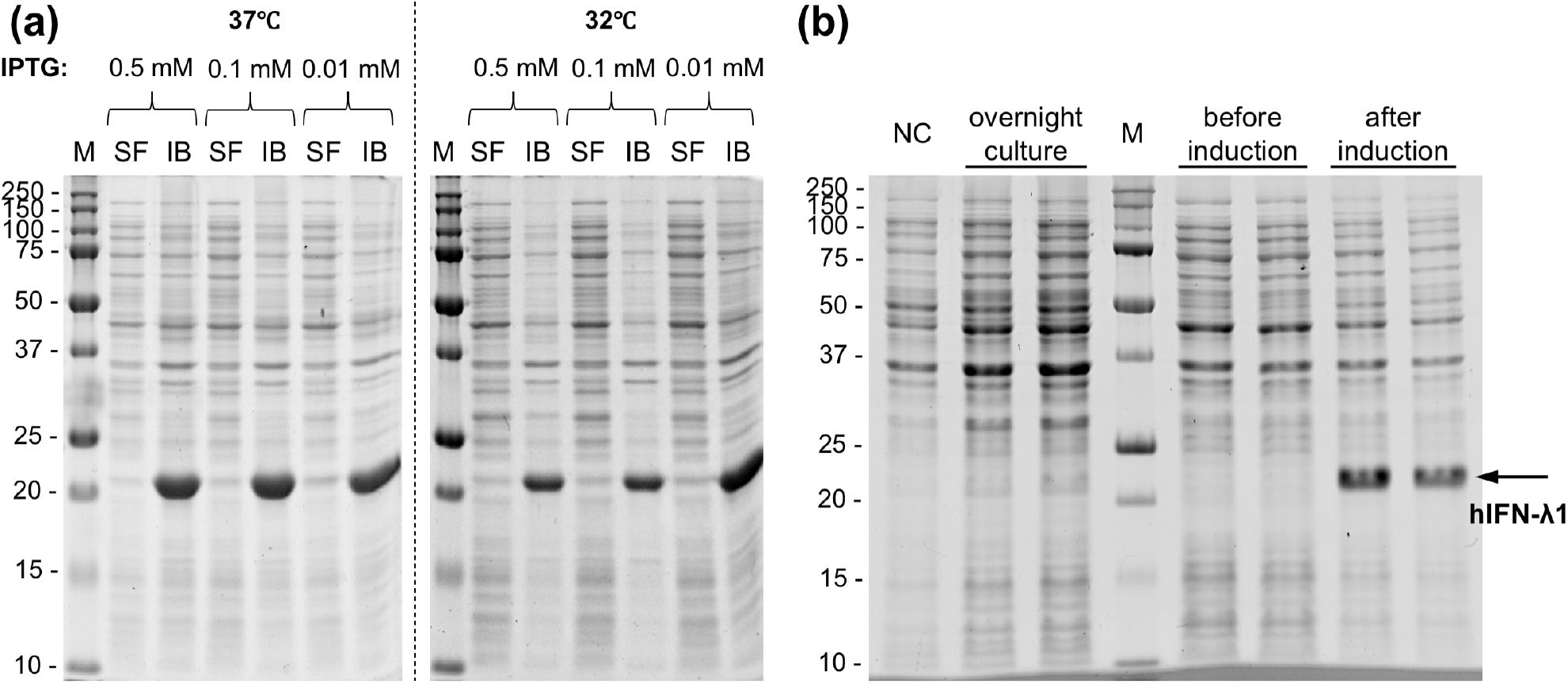
Electrophoresis of lysate fractions from the hIFN-λ_1_ producing strain (*E. Coli* BL-21). **(a)** Induction testing. The x-axes show IPTG concentration used for induction. Incubation temperatures are shown (left 37°C, right 32°C). SF – soluble fraction; IB – inclusion bodies. **(b)** Protein production. ‘Overnight culture’ - overnight culture, subcultured for production; ‘Before induction’ - cell lysate before addition of inducer; and ‘After induction’ - cell lysate, 3 hours after addition of inducer. NC – negative control (*E. Coli* BL-21 without plasmid).

Under the conditions tested, the target protein was expressed in the form of inclusion bodies. According to the literature [5,10,16], this is a common occurrence with production of recombinant interferons in *E. Coli*. It should be noted that the hIFN-λ_1_ gene was also cloned into the pet22b+vector, at the MscI and XhoI sites, in order to create a fusion protein not only with a histidine tag at the C-terminus, but also with pelB sequence at the N-terminus. The pelB sequence encodes a signal peptide, potentially contributing to periplasmic protein localization. However, even in the presence of the signal peptide, recombinant protein was localized in inclusion bodies (data not shown).

Since cultivation conditions and the presence of pelB sequence did not increase the amount of soluble protein, production of hIFN-λ_1_ was carried out using: pet302 NT-His plasmid; LB medium with ampicillin (100 μg/ml); 37°C incubation; and platform agitation (250 rpm). When conditions were met (culture OD_600_ = 0.7 arbitrary units), IPTG was added (0.05 mM final concentration), and cultivation with induction was continued for 3 more hours (Fig. 2b).

### 3.3. Protein purification and refolding

Primary protein purification was carried out by metal affinity chromatography under denaturing conditions. This purification method, with on-column refolding of hIFN-λ_1_ described earlier [10], showed low yield of refolded protein (less then 5 mg of correctly-folded protein per liter of culture). Therefore, it was decided to perform the first purification step under denaturing conditions, followed by refolding of the partially purified protein. Ni-Sepharose 6FF resin (prepacked HisTrap FF Crude columns) was chosen because of its high dynamic binding capacity (about 35 mg of hIFN-λ_1_ per 1 ml of sorbent) and the possibility of further scale-up. The chromatogram of the first purification step is shown in Figure 3a.

**Figure 3.**
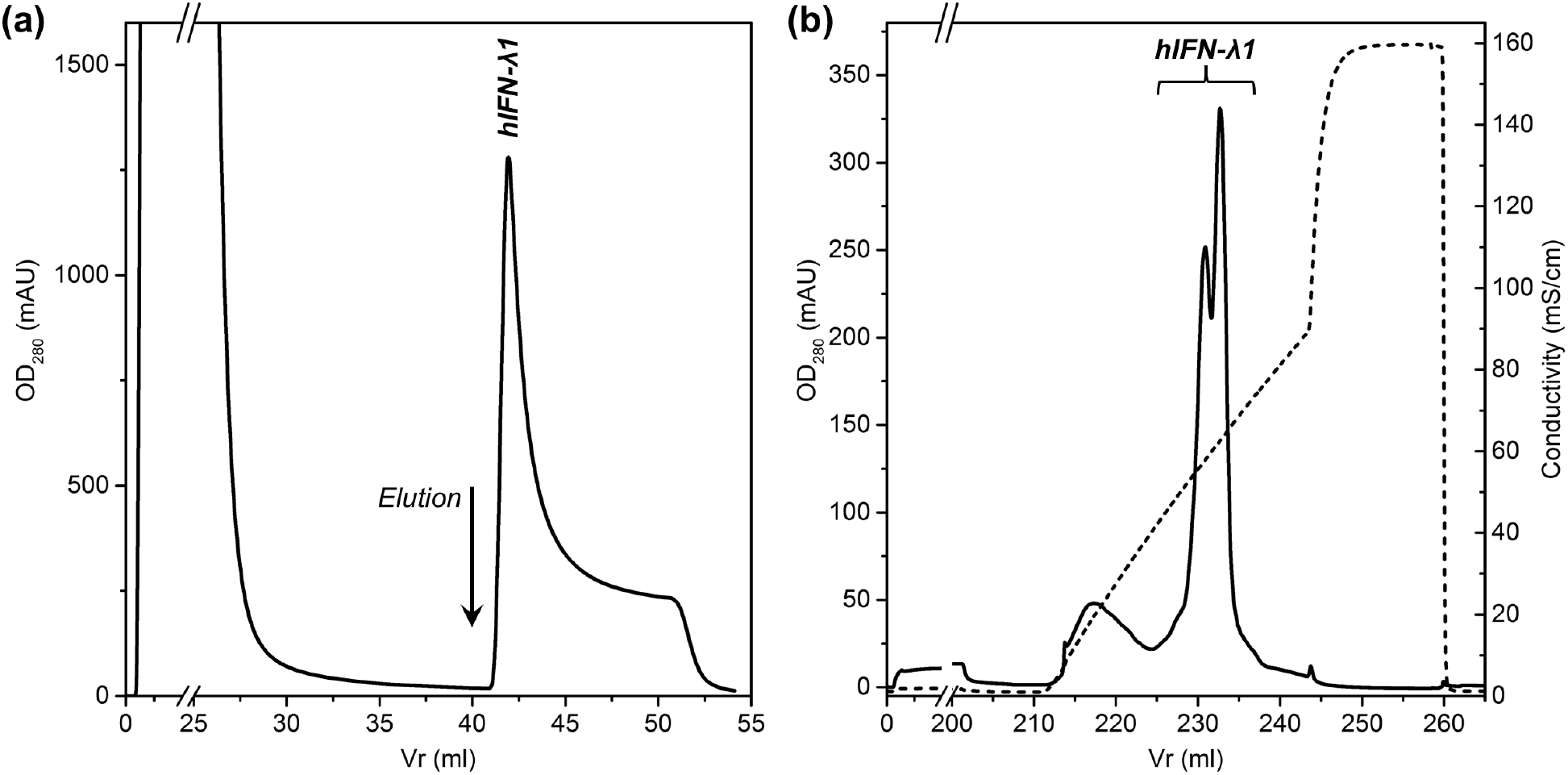
**(a) Primary purification of recombinant hIFN-λ_1_ by IMAC.** Absorbance at 280 nm is shown in blue; elution buffer concentration is shown in pink. **(b) Final purification of recombinant hIFN-λ_1_ by cation exchange chromatography**. Blue indicates absorption at 280 nm; red indicates conductivity of the solution.

Two refolding buffers with different pH (tris-HCl, pH 7.5; sodium acetate, pH 5.5), with and without refolding additives, were selected for screening. Denatured recombinant protein samples (at a concentration of 2 mg/ml) were added to pre-cooled refolding buffer (to final concentration of 100 μg/ml) and mixed by pipetting, followed by incubation for 24 hours at ^+^4°C. Precipitated protein was removed by centrifugation for 1 hour at 20,000 g. Supernatants were collected and dialyzed against phosphate buffered saline for 24 hours, followed by filtration through a PES syringe filter (0.2 μm pore size). Protein concentrations were determined by the Lowry method. For estimation of relative ‘soluble protein yield’ of refolding variations, 100 μg/ml was designated as 100%. The results of the experiment are shown in Table 1.

**Table 1.** Soluble protein yield in various refolding buffers.

| Refolding Buffer | Soluble Protein Yield, % |
| --- | --- |
| (A) 20 mM tris-HCl, pH 7.5 | 42 |
| (B) 20 mM tris-HCl, 1 M L-arginine, pH 7.5 | 55 |
| (C) 20 mM tris-HCl, 20% glycerol, pH 7.5 | 80 |
| (D) 20 mM tris-HCl, 5% trehalose, pH 7.5 | 40 |
| (E) 20 mM tris-HCl, 0.1% Tween-20, pH 7.5 | 89 |
| (F) 10 mM sodium acetate, pH 5.5 | 15 |
| (G) 10 mM sodium acetate, 5% trehalose, pH 5.5 | 12 |

The highest yields of recombinant hIFN-λ_1_ in soluble form were seen with two specific refolding buffers: Buffer C (80%); and Buffer E (89%). The proposed refolding method provides a significantly higher yield, takes less time, and is more economical in comparison with the stepwise dialysis method proposed [5] for IFN-λ_3_, another type III interferon. Conductivity of the protein solution after refolding was about 5 mS/cm; therefore, it does not interfere with subsequent ion-exchange chromatography. There is thus no need for dialysis or diafiltration of the refolded protein solution. It is possible to proceed directly to the final purification step.

The isoelectric point of the hIFN-λ_1_ recombinant protein was 9.08 according to the ProtParam Tool [17]. Thus, hIFN-λ_1_ in refolding solution is positively charged, so cation exchange chromatography was chosen as a final purification step. With comparison of different cation exchange resins, the best results (maximum protein yield, minimal impurities) were noted (data not shown) with resins featuring particle size of 15 μm or less, such as Source 15S (GE Healthcare) and ENrich S (Bio-Rad). Source 15S was selected because of higher flow rates and availability for purchase in bulk (for potential process scale-up later). The chromatogram of the final protein purification step is shown in Figure 3b. The target protein eluted in the conductivity range from 45 to 65 mS/cm. It should be noted that hIFN-λ_1_ elutes as a double peak, which may indicate the presence of two different conformational variants.

The method makes it possible to obtain up to 70 mg of recombinant hIFN-λ_1_ from 1 L of cell culture (∼5.2 g wet weight). Expression in the form of inclusion bodies protects the protein from proteolysis during cultivation. Simple and effective refolding provides a high yield of native-form protein with proven biological activity.

### 3.4. Product analysis

#### 3.4.1 SDS-PAGE and MALDI-TOF

The purity of the recombinant protein was analyzed by SDS-PAGE. Figure 4 shows all steps of target protein purification.

**Figure 4.**
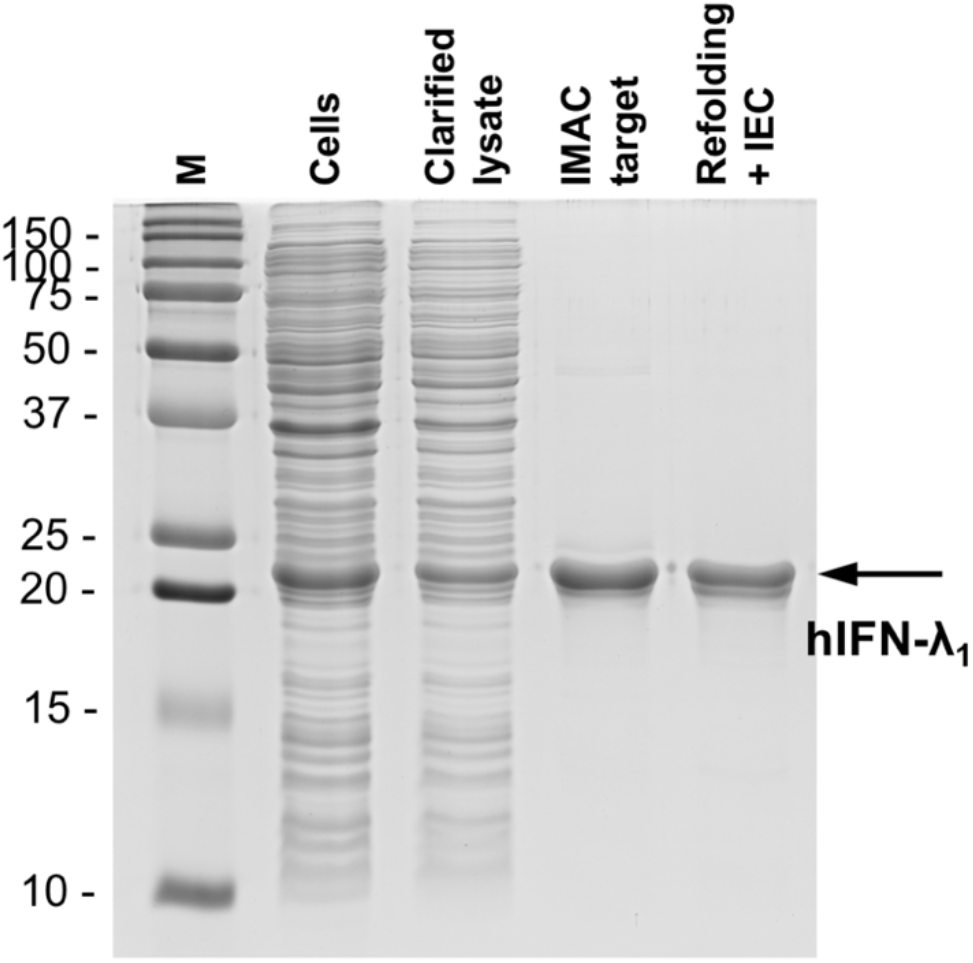
PAGE of 6His-hIFN-λ_1_-containing samples at various stages. Key: M – marker (Precision Plus mol. weight standard); ‘Cells’ – whole cell suspension; ‘IMAC target’ – target peak from metal-affinity column; ‘Refolding + IEC’ – material after refolding followed by cation exchange chromatography. Coomassie staining.

The presence of a band with an electrophoretic mobility of about 21 kDa (indicated by an arrow) indicates selective isolation of the protein from the protein mixture present in the cell culture.

Confirmation of recombinant protein amino acid sequence was carried out using MALDI-TOF mass spectrometry after enzymatic hydrolysis of ‘protein in gel’ with trypsin. In the area marked with an arrow (Fig. 4), the 6xHis-hIFN-λ_1_ recombinant protein was reliably identified (Score/threshold was 156/76, sequence coverage 70%). Additional hydrolysis with chymotrypsin extends the sequence coverage to 85%. Protein sequences found in mass spectra are highlighted in Figure 1b. Thus, the obtained protein was reliably identified as human interferon lambda-1 (6xHis-hIFN-λ_1_).

#### 3.4.3. ELISA

Binding of the recombinant protein with anti-hIFNλ_1_ monoclonal antibodies was shown by sandwich ELISA. The commercial ‘Human IL-29/IL-28B (IFN-lambda 1/3) DuoSet ELISA’ kit (DY1598B, R&D Systems) was used (Fig. 5).

**Figure 5.**
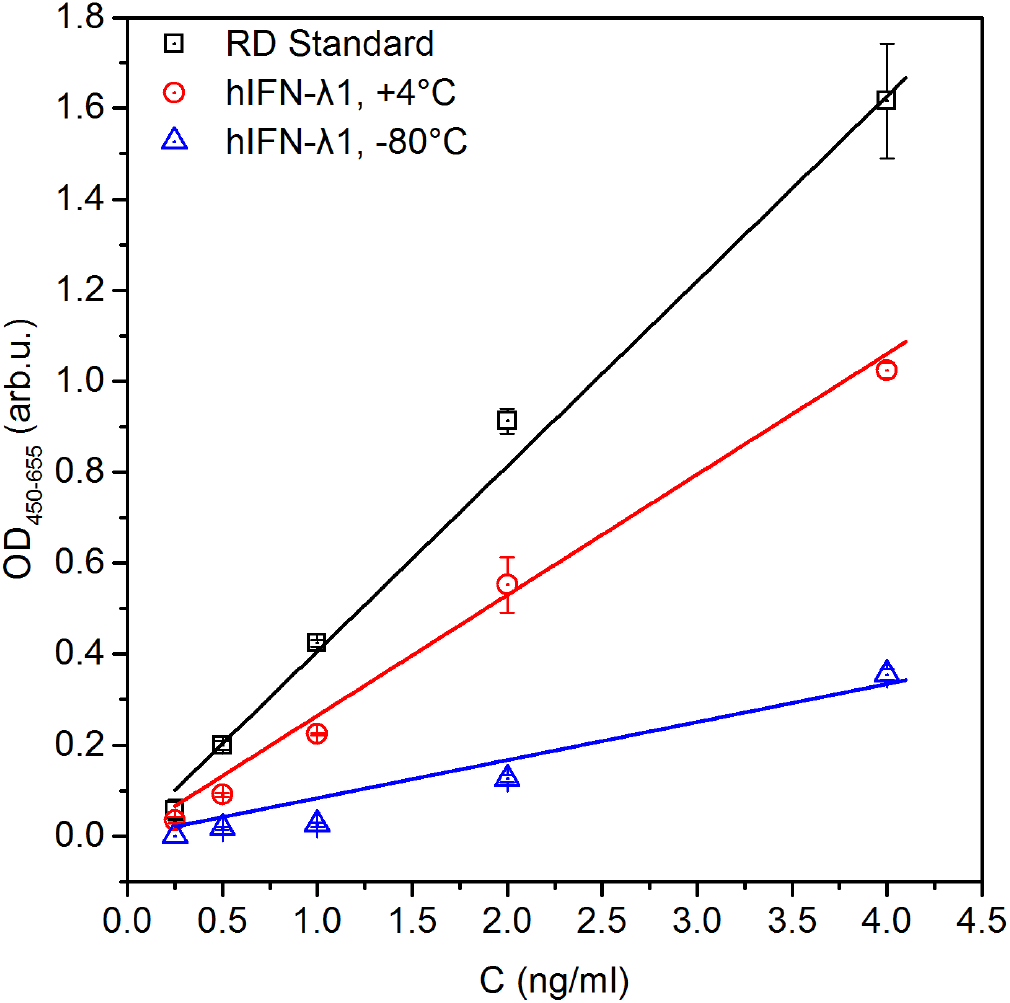
Comparison of ELISA signal from recombinant hIFN-λ_1_ stored in different conditions. In addition to the two storage conditions (+4°C and -80°C), a commercial standard is shown from the ‘Human IL-29/IL-28B kit (IFN-lambda 1/3) DuoSet’ ELISA kit (DY1598B, R&D Systems).

For each concentration, the average optical absorbance (OD_450_-OD_655_) is indicated in relative units. Concentrations of hIFN-λ_1_ were determined by the Lowry method. These results show that a constant ^+^4°C temperature is the preferred storage condition for recombinant hIFN-λ_1_ protein, not ^-^ 80°C followed by thawing. Increasing the stability of the protein product is a priority goal for further improvement of this method. Ideally, this would minimize changes associated with one or more ‘freeze-thaw’ cycles.

### 3.5. Confirmation of biological activity

#### 3.5.1. Limulus amebocyte lysate testing

In order to exclude the possibility of stimulation of A549 cells by bacterial lipopolysaccharides, the endotoxin level was determined by LAL test. The results show that recombinant hIFN-λ_1_ solution contains endotoxin at a concentration of about 25 EU/ml, which corresponds to no more than 5 ng/ml of endotoxin [18]. The concentration of the analyzed sample was 620 μg/ml. Consequently, there was less than 0.05 EU endotoxin per μg of protein, which is in line with commercial analogues on the market [19]. Thus, endotoxins present in the solution should not significantly affect the immune response of cells. Moreover, it has been shown that lipopolysaccharides do not induce the production of either IFN-α/β or IFN-λ in respiratory epithelial cells [14,20]. The A549 cells used here are also a respiratory epithelial line, and lipopolysaccharide-induced MxA expression is not expected

#### 3.5.2 Real-time PCR

The biological activity of hIFN-λ_1_ was shown by RT-PCR. Increased MxA expression was shown in A549 cells treated for 10 hours with hIFN-λ_1_ (10 ng/ml) compared to control cells (Fig. 6). The level of MxA expression increased by more than 20-fold. Since MxA is an interferon-stimulated gene, increased expression can be considered reliable evidence of the biological activity of the obtained hIFN-λ_1_ [14].

**Figure 6.**
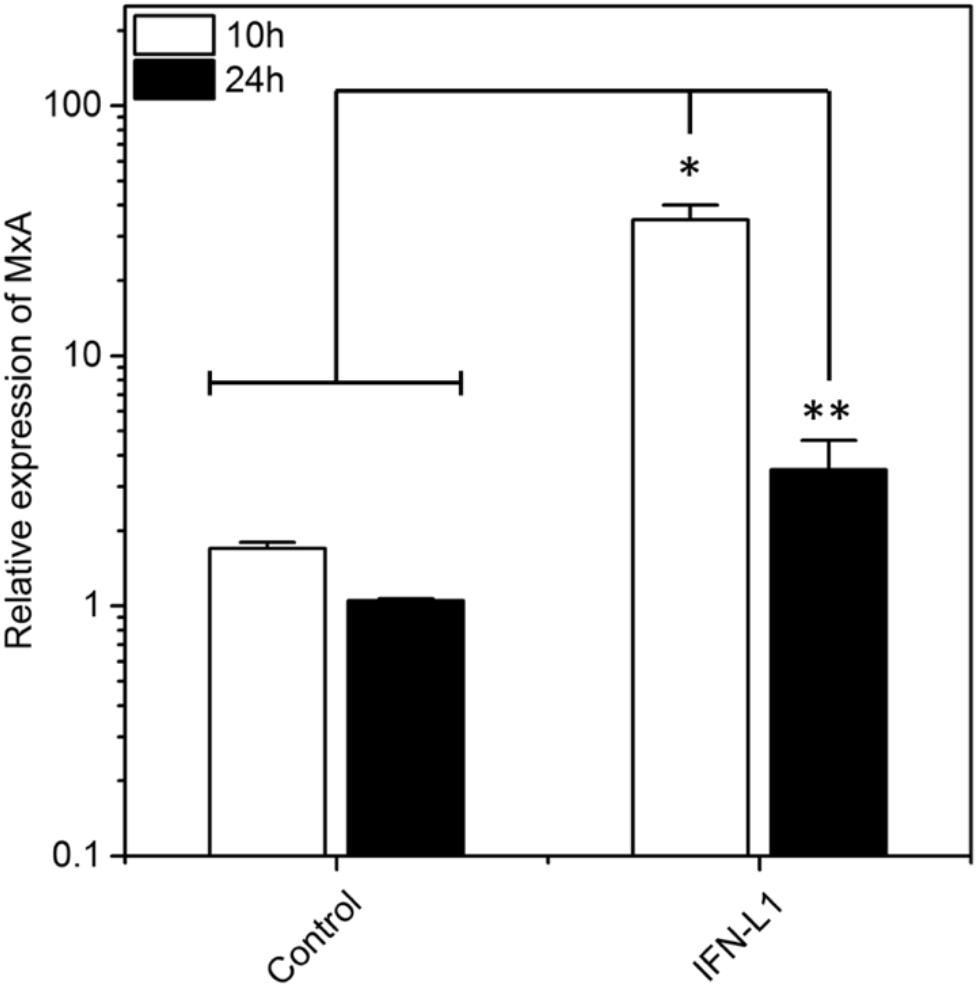
Change in MxA expression in A549 cells treated with hIFN-λ_1_ (10 ng/ml) for 10 or 24 hours during incubation. The y-axis shows relative MxA expression; the x-axis shows cell treatment conditions. The differences in MxA-2 mRNA expression between the control group at 10 h and the control group at 24 h were insignificant (p = 0.3333). T-test was performed using the nonparametric Mann-Whitney test. * p-value = 0.0286; ** p-value = 0.0159.

## AUTHOR CONTRIBUTIONS

were as follows: A.S. — methodology, data collection, data analysis, writing of manuscript; A.L., N.Y., M.P. — data collection, data analysis, writing of manuscript; T.K., A.T., P.N. — data collection, data analysis; Y.Z. — data collection, data analysis, writing of manuscript, visualization; E.R. — writing, review and editing of manuscript; M.G., A.V. — resources, supervision.

## ACKNOWLEDGMENTS

This work was supported by a Russian State Assignment for fundamental research (0784-2020-0023).

## CONCLUSION

In this work, we developed an efficient and potentially-scalable method for refolding and purification of recombinant human hIFN-λ_1_. The purified protein was confirmed to feature the expected biological activity. The developed method provides a high yield of protein with low impurity content, such as lipopolysaccharide. It is therefore potentially suitable for isolation of type III interferons not only for research and diagnostic applications, but also for therapeutic use.

